# Sex overrides mutation in *Escherichia coli* colonizing the gut

**DOI:** 10.1101/384875

**Authors:** N. Frazão, A. Sousa, M. Lässig, I. Gordo

## Abstract

Bacteria evolve by mutation accumulation in laboratory experiments, but the tempo and mode of evolution in natural environments are largely unknown. Here we show, by experimental evolution of *E. coli* in the mouse gut, that the ecology of the gut controls bacterial evolution. If a resident *E. coli* strain is present in the gut, an invading strain evolves by rapid horizontal gene transfer; this mode precedes and outweighs evolution by point mutations. An epidemic infection by two phages drives gene uptake and produces multiple co-existing lineages of phage-carrying (lysogenic) bacteria. A minimal dynamical model explains the temporal pattern of phage epidemics and their complex evolutionary outcome as generic effects of phage-mediated selection. We conclude that phages are an important eco-evolutionary driving force – they accelerate evolution and promote genetic diversity of bacteria.

**One Sentence Summary:** Bacteriophages drive rapid evolution in the gut.

## Main Text

The human gut harbors a densely populated microbial ecosystem, whose most abundant members are non-pathogenic bacteria. These commensals help in nutrient and drug metabolism and defense against pathogens, thus contributing to host health (*1*, *2*). Intra-species variation of commensals can also be critical for host-microbiota homeostasis (*3*). *Escherichia coli* is a common colonizer of the human intestine but also a potential pathogen. Comparative genomics studies show that *E. coli* evolves by mutation and recombination, *i.e.* bacterial sex, by which genetic material is shared between organisms that are not in a parent–offspring relationship (*4*, *5*). Horizontal gene transfer (HGT) has been recognized as a key factor in the long-term genome evolution of *E. coli*, but how much sequence divergence HGT generates compared to mutations is highly debated (*6*). On shorter time scales, the evolutionary speed and mode of *E. coli*, as well as their relative contributions to ecological adaptations, remain largely unknown. The mammalian gut is expected to represent a hotspot for HGT, but direct measures of the within-host HGT rate are scarce (*3*). The evolution of *E. coli* in the gut has been studied in mouse models, which are a classical system for *E. coli* physiology (*7*). In antibiotic-treated mice, mutations are the dominant mode of molecular evolution and HGT has not been reported (*8*, *9*). However, the antibiotic treatment in the standard experimental model can distort the tempo and mode of bacterial evolution in the gut, because it leads to a reduction in gut microbiota density and diversity (*10*), a possible depletion of *E. coli* competitors, and mutagenic effects.

To unmask the true evolutionary potential of commensal *E. coli*, we developed a new gut colonization model with a pre-adapted lineage (Table S1) that can grow in gut ecosystems of different diversity (*11*). In this model, an invading *E. coli* strain is able to colonize mice after a small perturbation of the host microbiota, and its evolution in the natural gut ecosystem can be followed (Fig. S1). Importantly, the gut microbiota of this system exhibit a significant variation across hosts (Fig. S5). In particular, prior to colonization, these microbiota contain a resident strain at variable loads between 10^3^ and 10^8^ CFU/g feces (Fig. 1A, Table S7). The resident *E. coli* carries two plasmids (with 68,935 bp and 108,557 bp) and is susceptible to a panel of 10 antibiotics including streptomycin (Table S1). The invading lineage belongs to *E. coli* phylogenetic group A and carries a yellow fluorescent marker (YFP) to allow its ready identification in fecal samples. This lineage is able to stably colonize the mouse gut for several months (data not shown).

**Fig. 1.**
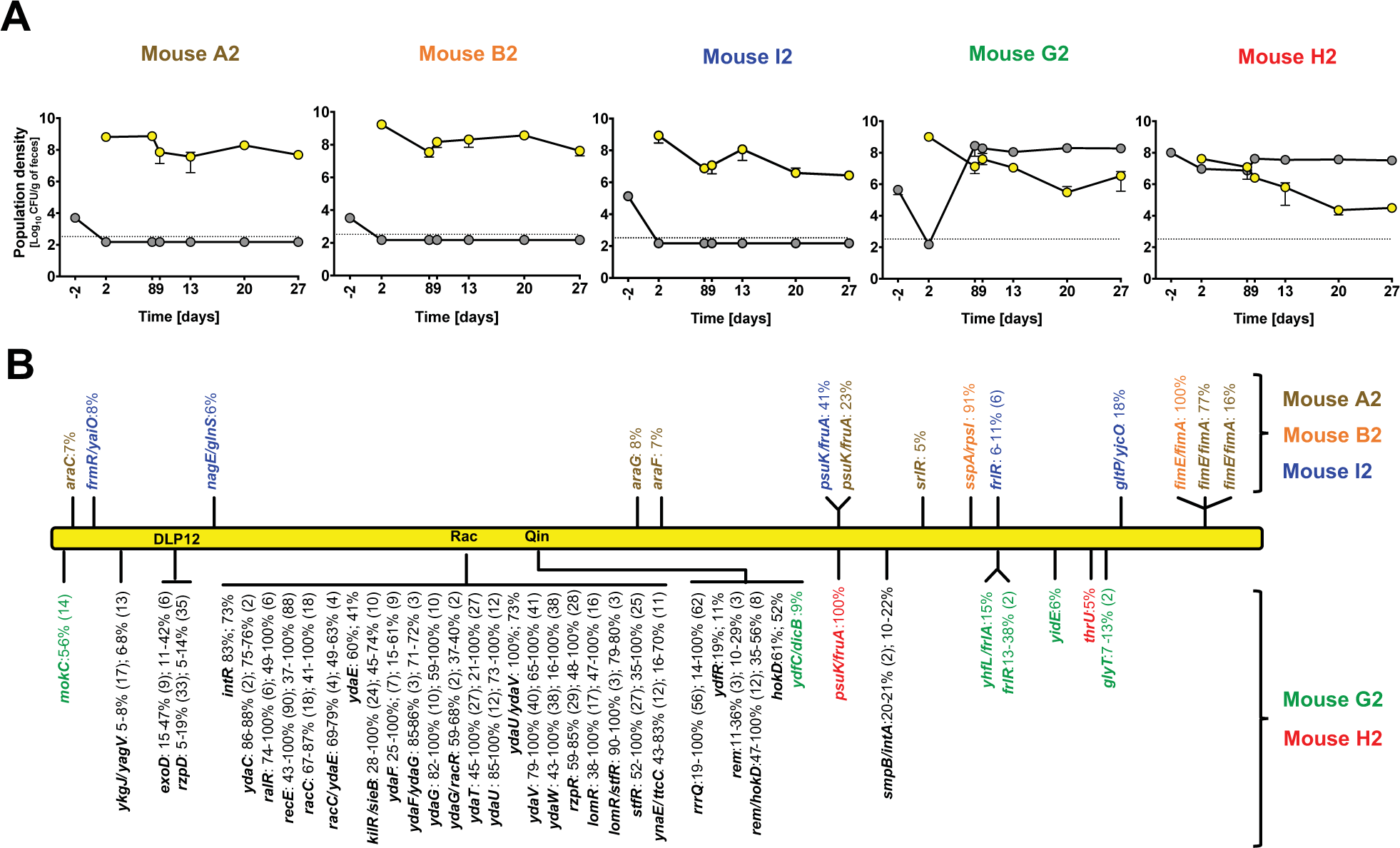
Replacement or coexistence of invader and resident *E. coli* strains during colonization. **(A)** Loads (log_10_ CFU/gram of feces) of invader (yellow circles) and resident (gray circles) *E. coli* in mice A2, B2, I2, G2 and H2 (error bars: 2SE; dotted line: detection limit). **(B)** Evolution in the invader lineage. Comparison of the evolved populations (day 27) with the ancestral clone by whole-genome sequencing: name of mutated gene or intergenic region, with mutation frequencies and number of distinct alleles in parentheses. Top: mutations in mice A2 (brown), B2 (orange), I2 (blue). Bottom: Mutations under coexistence; mice G2 (green), H2 (red). Genetic changes observed mainly within cryptic *E. coli* prophage (DLP12, Rac and Qin) sequences, with mutational parallelism in the sampled clones shown in black.

We colonized five mice with an invader *E. coli* strain and followed the population dynamics in the gut for one month after colonization (Fig. 1A). We found two distinct patterns. In three mice, the resident strain is lost and the invader settles to a stable population density (Fig. 1A, three leftmost panels). In the other two mice, the colonization leads to a stable coexistence of both strains, in which the resident regains the majority of the combined *E. coli* population (Fig. 1A, two rightmost panels). These outcomes of colonization can be linked to the initial load of the resident strain (Fig. 1A, Table S7): replacement was observed in mice with lower initial loads of the resident (10^3^-10^5^ CFU/g feces, mice A2, B2, I2); coexistence occurred in mice with a higher initial load (10^5^-10^8^ CFU/g feces, mice G2, H2). The combined load of resident and invader is maintained at an average of 10^8^ CFU/g feces across all mice, which we associate with the stable ecological niche size of the *E. coli* species (see Supplementary Text, Fig. S4, Table S7).

Strikingly, the two patterns of colonization are associated with drastic differences in the invader’s mode of evolution. Fig. 1B shows the results of whole-genome sequencing of the invader (pools of YFP clones) 27 days after colonization. When the resident lineage is absent, the invader evolves by successive accumulation of mutations, as seen before (*8*, *9*); when the two lineages coexist, we find a much higher rate of evolution marked by a vastly larger number of sequence changes in the same period, including numerous adjacent changes in known lambdoid defective prophage genes and sequence reads with no identity with the ancestor (Fig. 1B, Table S5). These changes reflect phage-driven HGT, as we show by sequencing and *de novo* assembly of an evolved invader-YFP clone sampled at day 27 from mouse G2 (Fig. 2A). This clone carries no detectable plasmids but displays an increased genome size (4.7 Mbp vs. 4.6 Mbp for the ancestral) due to the acquisition of two new prophages (Fig. S6 and Table S6). One of the prophages, comprising a new sequence of 49,357 bp, replaced the defective Rac prophage of the ancestral. As no signs of homologous recombination were detected in this clone, the data suggest that the invader YFP lineage exchanged its ancestral cryptic Rac prophage for a new Rac-like prophage acquired from the resident clone; we named this prophage KingRac. In addition, the invader acquired a new sequence segment inserted at the *ssrA* core gene (Fig. S6); we identified this sequence (47,407 bp) as a complete prophage and named it Nef (Fig. 2A).

**Fig. 2.**
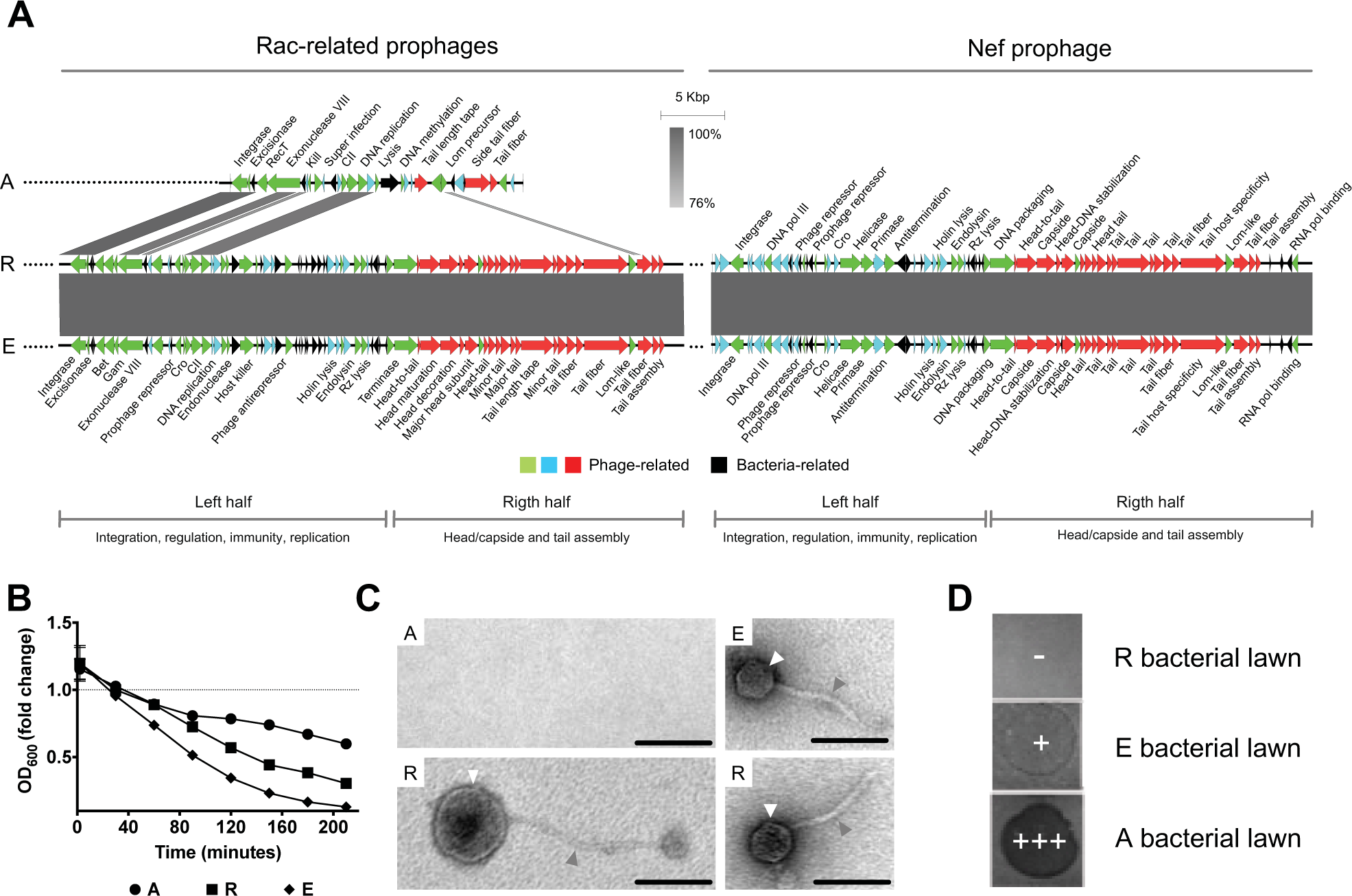
Phage-mediated horizontal gene transfer (HGT) causes evolution of invader *E. coli*. **(A)** Sequence alignment of prophage genomes present in the ancestral invader (A), the evolved invader (E), and the resident (R) *E. coli*. Two new prophages were found in the genome of the evolved clone: KingRac, a Rac-related prophage and Nef, a new prophage of unknown family. Arrows indicate coding sequences (CDS) found in the prophages (green, blue and red: phage-related CDS, black: bacterial CDS). **(B)** Phage induction assay. Population decline (fold change) under mitomycin C-induced bacterial lysis for *E. coli* clones A, R and E grown in LB media. **(C)** Electron micrographs of phage particles presenting two different morphologies. White arrows point to the phage head/capside, gray arrows point to the phage tail. A, E and R clones were induced with mitomycin C to produce phage particles. Bar = 100 nm. **(D)** Phage infection/lysis assay. Drops (10 µL) of phage-containing supernatant (phage lysate) produced by clone E were applied to growing bacterial lawns of R, E and A clones. Lysis pattern: -, no lysis; +, mild lysis; +++, strong lysis.

We used several methods to test whether the acquired prophages are active. In a phage induction assay (*12*), we found increased lysis of the evolved invader YFP clone relative to the ancestral (Fig. 2B), indicating that the acquired prophages can form active phage particles. Transmission electron microscopy of lysates showed that phage particles were produced by induction in the evolved and resident clones, but not in the ancestral clone (Fig. 2C, panel A, Fig. S7, Table S11). The evolved clone produced phage particles with an icosahedral head of 54 (±0.88, 2SE) nm and a tail length of 138 (±6.5, 2SE) nm (Fig. 2C, panel E), which were identical to those produced by the resident *E. coli* clone (Fig. 2C, panel R). The resident produced two morphologically distinct particles: one similar to phage lambda and similar to that detected in the evolved invader, and another displaying a spherical morphology (head 99,6 nm; tail 256,8 nm) (Fig. 2C, panel R). We tested the infective power of the acquired phages by an infection/lysis assay (see Supplementary Materials). We found a clear ranking of strains: the resident strain is immune, the evolved invader is moderately susceptible, and the ancestral invader is highly susceptible to infection with cell lysis (Fig 2D). Together, our results show that active prophages can readily succeed cryptic phages in multi-strain microbiota. The observed differences in induction and infection rates (Fig. 2BD) are key components of our phage-mediated selection model; see equation (1) below.

Next, we inferred the relative contributions and the temporal order of HGT and point mutations. Under conditions of strain coexistence, the invader populations accumulate only few mutations that reach a frequency ≥ 5% (2 in H2, 4 in G2) over one month. These numbers are comparable to the mutation accumulation in the other mice (2-7 in A2, B2, I2) (Fig. 1B). The faster temporal dynamics of HGT relative to *de novo* adaptive mutations at the *frlR* gene and *psuK*/*fruA* intergenic region was inferred by amplicon sequencing, time-resolved target PCR and whole-genome sequencing of pools of clones from mice G2 and H2 (Fig. 3AB). In both cases, the invader incurred a rapid succession of two phage infections driving HGT (lysogenization). The first, Nef-driven HGT event occurred just two days after colonization; this was followed by the integration of KingRac (Fig. 3AB). We conclude that carrying Nef does not confer full immunity to infection by KingRac; *i.e.*, Nef and KingRac can be seen as the first and second line of the resident’s defense against the invader. Moreover, *de novo* adaptive mutations at the *frlR* locus of the invader genome spread only after the initial Nef-driven HGT (Fig. 3A, Table S15).

**Fig. 3.**
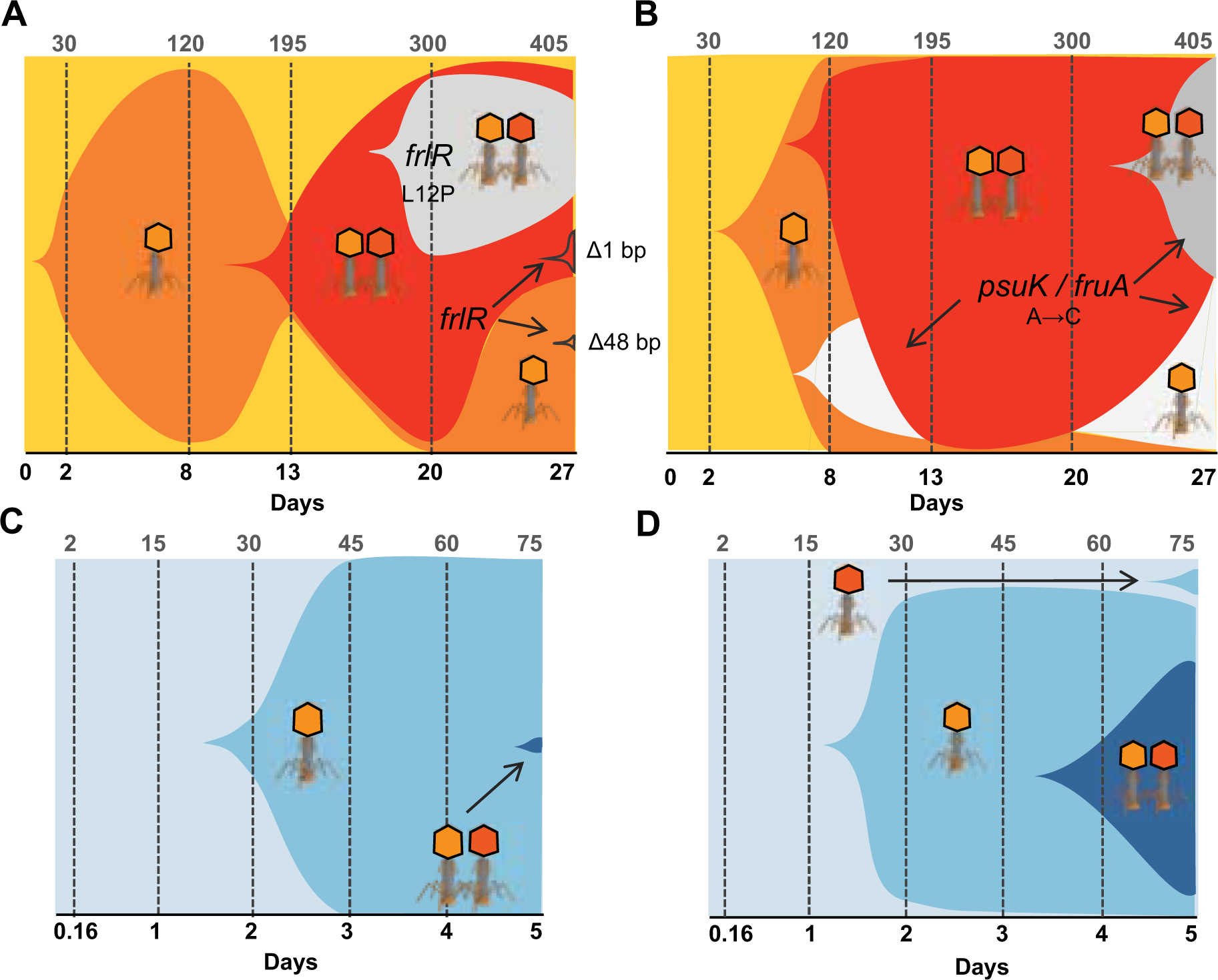
Phage-mediated HGT precedes evolution by point mutations. Muller plots representing the dynamics of the phage-mediated HGT and mutation processes in the invader *E. coli* population. Shaded areas indicate the frequency of each genotype; phage symbols represent single or double lysogens. Colonization time is given in days (lower axis) and estimated number of generations (upper axis). **(A, B)** Evolution experiments: ancestral clone (orange), lysogenization by Nef (dark orange) followed by KingRac (red), and *frlR* (in A, grey) or *psuK*/*fruA* (in B, grey) mutations. **(C, D)** Co-colonization experiments: ancestral clone (light blue), lysogenization by Nef and KingRac (darker blue shades).

To evaluate the reproducibility of the observed HGT dynamics, we co-colonized a new cohort of mice with both the evolved YFP lysogenized clone (extracted at day 27 of the previous experiment) and the ancestral invader clone expressing a cyan fluorescent protein (CFP) marker (Table S1). Stable gut colonization of both clones was observed within five days (Table S8). The ancestral (CFP) clone incurred a Nef-driven HGT event within two days in all mice, and a KingRac-driven HGT event always occurred after the Nef event (Fig. 3CD, Table S10).

The Muller plots of Fig. 3 depict the speed and mode of evolutionary change at both the core and mobile genomes of the invader strain. They show a fast and repeatable pattern of phage-driven HGT. Evolution by HGT can reach the time scale of ecological change (see Fig 1A) – it precedes and outweighs evolution by point mutations throughout the duration of the experiment (Fig. 3AB). The HGT and lysogenization dynamics found here in commensal *E. coli* are faster than those previously observed in a murine diarrhea model of Salmonella infection (*13*) and in a rat model of lung infection by *Pseudomonas aeruginosa* (*14*), suggesting an important rolefor HGT not only in disease but also in health. Importantly, the observed HGT did not eliminate neutral genetic diversity in the core genome: Nef and the genetically-linked new bacterial genes swept close to fixation, while a polymorphism of CFP/YFP markers was maintained in the invader population (Fig. 3CD, Table S8). Hence, phage-mediated HGT can lead to the spread of new genes, while keeping high genetic diversity in the core genome.

The bacterial evolution of Fig. 3 also reveals a striking pattern of increasing diversity, which is marked by coexistence of invader and resident strains. This pattern is very different from the prevalent mode of evolution in laboratory experiments, which is characterized by rapid fixation of adaptive point mutations. However, the presence of phages introduces selection depending on the phage load, which provides a parsimonious explanation of the eco-evolutionary diversity. Differences in induction and infection rates between co-existing strains, as observed in our system (Fig. 2BD), contribute to this selection. Specifically, the fitness difference (selection coefficient) between two strains,

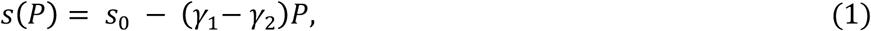

depends on the phage population density, *P*, and the fitness difference in the low-page limit, *s*_0_ = (*f*_1_ − *f*_2_) − (*δ*_1_ − *δ*_2_). Here *f*_i_ denotes the background fitness of strain *i* without induction or phages, *δ*_i_ is the fitness cost of induction, and *γ*_i_ is the fitness cost of infection per unit of phage, which is proportional to the infection rate (*i* = 1,2). For example, a more susceptible strain 1 competing with a more inducible strain 2 (*γ*_1_ > *γ*_2_, *δ*_2_ > *δ*_1_) can have a selective advantage at low phage levels but a disadvantage at high levels. In turn, the phage load at a given time depends on the population sizes of phage-producing bacteria in the recent past. This selective feedback can lead to stable coexistence of bacterial clades with different infection rates.

To show that the pattern observed in Fig. 3 is consistent with phage-mediated selection, we use a minimal eco-evolutionary model: a susceptible and an inducible (lysogenic) strain compete in a single ecological niche of a given carrying capacity; see Methods and refs. (*15*, *16*). Phages are produced by infection and by induction, and they are cleared with a constant rate. The model population dynamics can take the form of an infection epidemic (Fig. 4A), with a rapid increase of phages, causing negative selection (*s* < 0) and a resulting depletion of susceptibles. In the second phase, low phage production and clearance lead to lower phage levels, positive selection (*s* > 0) and a rebound of susceptibles. This pattern of time-dependent selection is observed during the first two weeks of colonization in mouse G2 (Fig. 3A). Variation of the initial conditions, for example of *P*, generates epidemics of different strength (Fig. S9), ranging from a rapid loss of susceptibles (as observed in Fig. 3BC) to a monotonic decline of susceptibles (as observed in Fig. 3D).

**Fig. 4.**
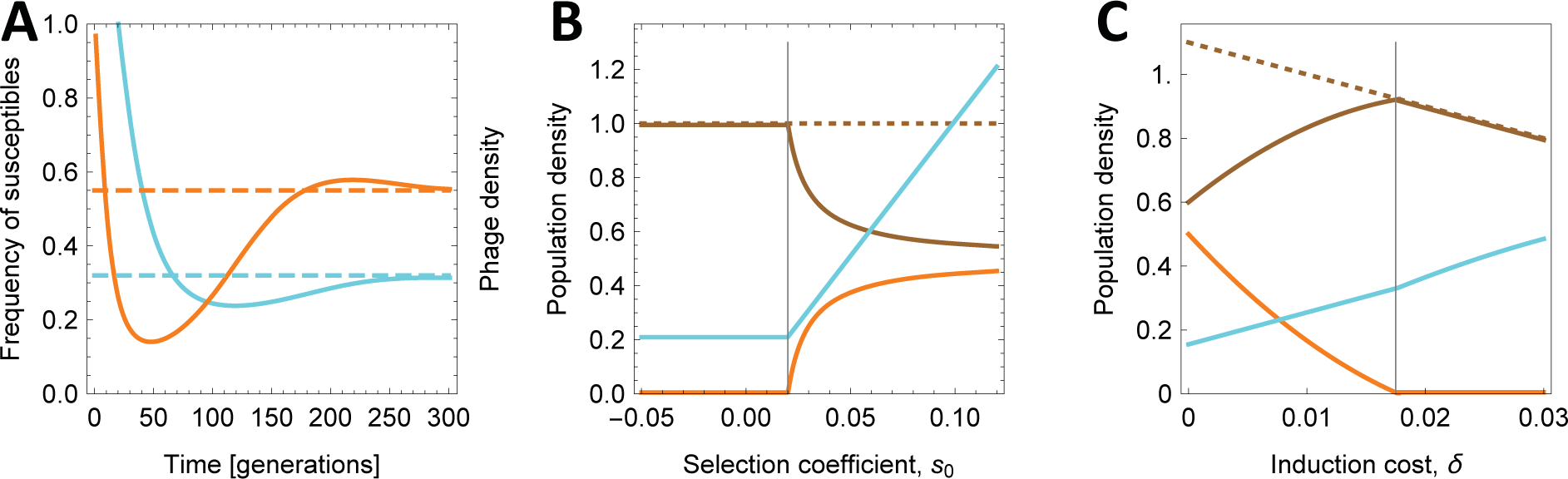
Phages generate epidemics and promote coexistence of bacterial strains. **(A)** Phage infection epidemics. The population frequency of susceptible (ancestral) bacteria in the invader population (orange) and the phage population density (cyan) are plotted against time (stable equilibrium values are marked by dashed lines). The epidemic pattern is marked by an initial decline of susceptibles at high phage levels, followed by a rebound at lower phage levels; see also Fig. S9. **(B)** Equilibrium population sizes (brown: inducible, resident strain; orange: susceptible, invader strain; dashed brown: total bacteria load; cyan: phage) are plotted against the selective difference at low phage density, *s*_0_. Beyond a threshold value 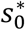 (vertical line), there is stable coexistence of two bacterial strains in the presence of phages. **(C)** The same equilibrium is plotted against the selective cost of induction, *δ*. Population size and fitness of the inducible strain (brown line) reach a maximum close to the onset of coexistence, *δ*^∗^(vertical line). Model parameters: infection cost *γ* = 0.1, induction cost *δ* = 0.01, lysogenization fraction *κ* = 0.5 (in A), background fitness *f*_*S*_ = 0.15, 0.125 (in A, C), *f*_*I*_ = 0.11, carrying capacity *c*^-1^ = 0.1, phage clearance rate *λ* = 0.05; definitions and model details are given in Materials and Methods.

Importantly, the model highlights how the complex outcome of the evolution and co-colonization experiments (Fig. 3) can arise. It shows that an ecosystem with phages can generate stable coexistence of strains even under conditions where one of the strains would always be displaced by the other in the absence of phages (see Methods). Coexistence in the presence of phages, as observed in our experiments, signals that the susceptible (invader) strain has a fitness advantage against the more inducible (resident) strain at low phage density, *s*_0_, that exceeds a threshold value 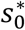 (Fig. 4B). A positive value of *s*_0_ is consistent with the observed difference in induction rates (Fig. 2B). Furthermore, plotting the equilibrium population sizes against the induction rate of the resident strain, δ, shows that the resident population size reaches a maximum at a value *δ*^∗^ close to the onset of coexistence (Fig. 4C). Rates close to *δ*^∗^ indicate the resident’s optimum investment in phage-mediated defense: they balance the cost of induction in the absence of competitors with its benefit as a defense mechanism against invaders (*17*, *18*).

In summary, this study demonstrates that phage-driven horizontal gene transfer can be a rapid and common mode of evolutionary change in gut microbiota. The underlying phage-mediated selection, by which susceptible strains can have an advantage at low phage levels and inducible lysogenic strains have an advantage at high phage levels, leads to competition warfare between strains. The resulting evolutionary dynamics is remarkably similar to resource competition observed in yeast populations, in which producers (cheaters) have an advantage at low (high) levels of a common good (an extracellular nutrient of yeast) (*19*). In our system, phages can be regarded as a “common load” that is produced by the distinct mechanisms of infection and induction but affects all bacteria within a population. To understand the long-term effects of these dynamics on microbiota strain variation remains an important challenge for future work (*20*). Assuming invasion, phage-mediated defense, and coexistence patterns observed experimentally as recurrent processes of natural *E. coli*, the model predicts that commensal strains should produce balanced, intermediate induction rates that allow coexistence with more susceptible strains. Thus, phage-mediated eco-evolutionary dynamics should promote bacterial diversity.

